# Punishment Risk Task: Monitoring anxiogenic states during goal directed actions in mice

**DOI:** 10.1101/2022.10.30.514458

**Authors:** Kyle E. Parker, Joel S. Arackal, Sarah C. Hunter, Carl Hammarsten, Jordan G. McCall

## Abstract

Canonical preclinical studies of anxiety-related behavioral states use exploration of novel spaces to test approach-avoidance conflicts such as the open field test, elevated plus maze, and light-dark box. However, these assays cannot evaluate complicated behaviors in which competing states of motivation result in anxiogenic behaviors. Furthermore, these assays can only test the approach-avoidance conflict once due to a reliance on spatial novelty. Here we demonstrate the punishment risk task (PRT) in male and female, group- and singly-housed mice, a model initially described in singly-housed male rats by Park and Moghaddam (2017). The task tests how probabilistic punishment affects reward-seeking behavior. In particular, it measures the delay to pursue a reward (sweetened food pellet) while the likelihood of punishment (foot shock) actively impinges reward-associated actions. Here, we found that mice show increased latency to respond to food reward cues in trials in which the probability of punishment is highest. Further, anxiolytic treatment with diazepam or propranolol block any increase in response latency, indicating the model’s potential to for study of anxiogenesis in mice. Elucidating how these competitive behavioral states are integral to adaptive behavior and change over time and experience to coordinate anxiogenesis should greatly benefit anxiety disorder research. Specifically, implementing this assay in mice will enable cell-type selective interrogation of these processes and further our understanding of the neural basis of anxiogenesis.

## Introduction

Animal models are key to understanding the etiology and neurobiology of neuropsychiatric disorders such as anxiety and depression. Unfortunately, the heterogeneity of anxiety disorders in humans combined with complex diagnostic criteria produces obstacles for modeling clinical conditions in animals[1,2]. In an effort to model human pathological anxiety in rodents, an extensive range of behavioral models have been established[3–10]. Many of these assays use simplistic approaches designed to evaluate animals’ natural responses to aversive stimuli or contrast their propensity to approach or avoid exploratory behavior or social interaction against the aversive properties of novel open, brightly lit, or elevated spaces (i.e., open-field, light-dark box, and elevated zero/plus mazes). These novelty-based physical exploration assays limit assessment of more complex behavioral states in which competing motivations can impede normal functioning and evolve over time. Parsing how these explicitly competing behavioral states are integral to adaptive behavior, predict susceptibility to stressors, and coordinate anxiogenesis will greatly benefit anxiety disorder research.

Recently, a rat behavioral model, initially described by Park and Moghaddam (2017)[11], assessed reward seeking behavior in context of increasing punishment probability. Here animals perform an instrumental action (i.e., nosepoke) on a fixed ratio-1 (FR1) schedule to receive a reward (i.e., sugar pellet). In the initial block of the task, animals perform a simple FR1 schedule. However, as each block passes, some trials are pseudorandomly punished with an electrical footshock. The probability of this punishment increases in both the second and third block. Despite this block-wise increase in punishment probability, the FR1 reward schedule remains the same across all three blocks. In this manner, animals display increased latency to perform the rewarded actions as the probability of punishment increases. This increase in punishment-induced latency suggests a transient state of hesitancy that dramatically affects performance during the session, particularly within the final block which has the highest probability of punishment. Diazepam, a well-established anxiolytic, significantly diminished this latency in within-subject trials compared to saline-treated sessions, demonstrating the predictive validity of the task as a model to induce and examine anxiogenic behavior. Importantly, the rats show distinctly slower reaction times, but not reward reaction times, as entries to procure the food are left unaffected. The implication is that punishment likely does not induce anhedonia or avolition but generates a behavioral state in which a small probability of punishment causes animals to be hesitant to pursue known reward-related actions. We hypothesize that this behavioral state is anxiogenic. If so, this model could be used to evoke an anxiogenic states, predict susceptibility to stressors, and test potential therapeutics.

Many experimental procedures evaluate anxiety states following exposure to unfamiliar aversive environments that can be elicited or exacerbated by a range of threatening stimuli such as predator exposure or pain[3,10,12]. Although baseline behaviors are presumably appropriate and adaptive for the present conditions in rodents, anxiety disorders in humans constitute maladaptive or pathological responses to the existing environment[1], suggesting that the ideal rodent behavioral assay could be tested repeatedly in the same animals. Here we have extensively adapted and tested the punishment risk task (PRT) introduced in rats[11] to demonstrate its validity in mice. Given the importance of housing and testing conditions on behavior[13–16], we have replicated this model in isolated mice and extended our testing to include group housing and foot-shock amplitude. Understanding performance in this task under different housing and punishment conditions is essential to determining how it can best be used going forward. The work we present here provides a foundation for more comprehensive interrogation of the neural circuitry underlying behaviors related to anxiety. Widespread use of this behavioral model in mice will enable critical studies with genetic tools to alter gene expression and enable *in vivo* imaging and neuronal manipulation during behavior. Because of the task-based nature of this repeatable behavioral model, important temporal dynamics of the underlying circuit function can be probed with this approach[11]. When fully-implemented across species, this assay could provide important translational insights to inform the development of novel treatment strategies for anxiety disorders.

## Methods

### Subjects

Male and female C57BL/6J (RRID:IMSR_JAX:000664) 3-6 month old mice were bred locally in a different building. Upon weaning, all animals were group-housed. Mice were transferred to the experimental facility and allowed 1 week of habituation. Animals were gently handled weekly (e.g., picked up by the base of the tail and placed on a scale). During PRT training and testing, mice were handled daily. Prior to any drug administration experiment, mice were briefly restrained by hand and lightly stroked on the abdomen by the experimenter once a day for three days. For pilot experimentation, male and female mice (n=14) were kept group-housed for the entirety of the experiment. For social isolation, mice were separated into singly-housed cages (n=16) or kept in their original group-housed cage (n=16). After 4 weeks of isolated or grouped housing, mice were tested for canonical behavioral phenotypes in OFT and EPM. One week later, mice were weighed and placed under a restricted diet to maintain 90% of free feeding bodyweight. Group-housed animals were placed into their respective cages and given equal weight pellets (2.5g for males, 2.0g for females) and observed to confirm consumption. Animals consume the pellets within 15 minutes. After one week, animals were trained for PRT. Mice had free access to water. Mice were maintained on a 12:12-hr light/dark cycle (lights on at 7:00 AM). Animals were weighed daily to confirm and maintain food restricted weight throughout the experiment. All procedures were approved by the Institutional Animal Care and Use Committee of Washington University (#20-0139), conformed to US National Institutes of Health guidelines, and the ARRIVE guidelines (Animal Research: Reporting of In Vivo Experiments) were followed as closely as possible.

## Behavior

### Open Field Test

OFT testing was performed in a sound attenuated behavior testing room as previously described[17,18]. Lighting was stabilized at ∼12 lux. Animals were allowed to habituate to the room for 1 hour prior to testing. Each animal was individually placed into a 2500 cm^2^ enclosure for 20 minutes. The center of the apparatus was defined as a square equal to 50% of the total area. Activity was video recorded via a Google Pixel 3 XL and videos analyzed using Ethovision XT 13 (Noldus Information Technology, Wageningen, The Netherlands). Distance traveled and time spent in the center and periphery zones of the apparatus were determined and averaged for each animal.

### Elevated Plus Maze

EPM testing was performed within a sound attenuated behavior testing room. Lighting was stabilized at ∼12 lux. Animals were allowed to habituate to the room for 1 hour prior to testing. Each animal was individually placed into a plus-shaped platform for 15 minutes. The apparatus is comprised of two open arms (25 x 5 x 0.5 cm) across from each other and perpendicular to two closed arms (25 x 5 x 16 cm) with a center platform (5 x 5 x 0.5 cm) all 50 cm above the floor. Activity was video recorded via a Google Pixel 3 XL and videos analyzed using Ethovision XT 13 (Noldus Information Technology, Wageningen, The Netherlands). Distance traveled and time spent in the open and closed arms was determined and averaged for each animal.

### Punishment Risk Task

The Punishment Risk Task (PRT) was performed within sound-attenuated boxes (Med Associates Inc., Fairfax, VT). Mice were trained to nosepoke on an FR1 schedule in response to a combined auditory (1s, 1000 Hz) and light cue inside the associated cue port to receive a sweetened pellet (Dustless Precision Pellet-F0071, Bio-Serv, Flemington, NJ). The light cue remained on until the mouse completed a nosepoke. Completion of nosepoke entry was determined by disruption of an infrared activity monitor located inside the nosepoke port. Immediately following completion of the nosepoke, the port light turned off and a house light illuminated the chamber and a sweetened food pellet was dispensed into the food hopper. Once the mouse entered the food hopper to retrieve the pellet, the house light turned off and a randomized intertrial interval of 8-13 seconds was initiated, followed by the next trial and cue presentation. The nosepoke port and food hopper were located on opposite walls of the operant chamber. The fixed ratio remained at one throughout all training and test sessions. Specifically, each mouse received one sweetened pellet after every completed nosepoke entry. Each mouse completed daily training sessions of 45 trials (60 minutes session duration, 10 days total) prior to the testing session. After training sessions, mice underwent a no-shock control test session, throughout which they performed 45 trials of nosepokes divided into three blocks separated by 2 minutes of darkness without any probability of punishment. Next, mice performed the final test that includes increased probability of punishment. The punishment contingencies were 0 for *Block 1*, 0.066 for *Block 2*, or 0.133 for *Block 3*. The punishment employed in this task is electrical foot shock (0.1 mA or 0.3 mA, 300 ms). Each block consisted of 15 trials with the associated contingency value assigned to each block in ascending order of probability. That is, in *Block 1* there was a 0% chance of foot shock, in *Block 2* there was a 6.66% chance of foot shock, and in *Block 3* there was a 13.33% chance of foot shock. Each block was separated by 2 minutes of darkness to help discern a change in block. In *Block 2*, mice were randomly assigned to received one shock upon nosepoke number 2, 3, or 4. In *Block 3* mice were randomly assigned to received one foot shock upon nosepoke number 2, 3, or 4 then a second foot shock upon nosepoke number 6, 7, 8, or 9. This pseudo-randomization of the shock presentation follows the shock parameters used in Park and Moghaddam (2017) [11]. As previously described[11], we used a maximum cutoff of 60 minutes for each block for all training and test sessions. Sessions that were terminated at this cutoff were excluded. The number of mice and session that were terminated was recorded. <u>Rationale for exclusion of mice</u> <u>for non-completion:</u> The expectation of this assay is that the experience of a punishment will induce an increased latency between cue and nosepoke. Because of this, for each mouse we expect the distribution of latencies across 15 trials in each block and across all three blocks to have a positive-skew. Essentially, we expect the later-recorded latency values to be higher than the earlier-recorded values. Likewise, we expect to see a shift upward in our data set, on a per-mouse basis. To better capture this aspect of the data in our summary statistic, we then use mean rather than median when aggregating our observations for each mouse. We aim to compare the collection of mean latency times for different groups of mice in different sexes, blocks, or treatments. To do this, it is important that these individual mean values be determined as uniformly as possible. Specifically, each mouse needs to complete the exact same number of trials so that our computations come from as alike of sample distributions as is reasonably controllable. Excluding mice that do not successfully complete all 15 trials in the 60-minute time limit per block ensures this equality of sample size for each mouse. We did not observe habituation to the foot shock as evidenced by lack of behavioral changes across sessions. Animals that did not reach training criteria or complete any trial block within 60 minutes were excluded. Throughout all complete sessions, nosepoke latency and reward latency was recorded. Nosepoke latency refers to time between auditory/light cue onset and completed nosepoke and reward latency refers to time between reward delivery and reward retrieval. PRT locomotor activity was captured via a Google Pixel 3 XL and videos analyzed using Ethovision XT 13 (Noldus Information Technology, Wageningen, The Netherlands).

## Drug Treatment

Diazepam (Sigma-Aldrich, St. Louis, MO, USA) and propranolol hydrochloride (Tocris, Bristol, UK) were diluted in 0.9% physiological saline and prepared on treatment days. For each drug, experimental conditions included three separate test sessions, each consisting of three blocks. Each mouse received an intraperitoneal (i.p) administration of saline for the first and last session and diazepam administration (2 mg/kg, i.p.) or propranolol (10 mg/kg, i.p.) for the second test session. Drug doses were selected based on previously published works examining anxiolysis in mice. These saline treatment tests were to determine the effects of injection and post treatment effects. Following treatment administration, each mouse was immediately returned to their homecage for 10 minutes prior to PRT. Each test session was separated by three days.

## Data Analysis & Statistics

Given the substantial effect previously observed in rats [11], we estimated the effect size to be 0.5 and used an *a priori* repeated measures ANOVA power analysis (0.95) (G*Power 3)[19] that indicated a required sample size of 12. In practice we typically began with a sample size of 15 as our mice are typically housed five mice per cage. Cages of mice were randomized to be group-or single-housed. Complete blinding with this strategy is not possible due to clear evidence as to whether a cage houses one or more mice. However, all behavioral data was automatically collected either from Med Associates code (Med Associates Inc., Fairfax, VT) or Ethovision XT 13 (Noldus Information Technology, Wageningen, The Netherlands). Given that, if less than 75% of animals did not reach the 45 successful operant response criterion for inclusion we would not meet the power requirements described above. Therefore, starting group sizes for approach-avoidance measures were n = 21 (group-housed) and n = 20 (singly-housed). Final group sizes for PRT testing were n = 13 (group-housed) and n = 15 (singly-housed). Pilot experimentation had an initial group size of n = 10. If an animal did not complete the PRT test, they were not included in analysis. All PRT data was collected via Med-Associates hardware and software or Ethovision XT 13 and were averaged and expressed as mean ± SEM. Each data point in the graphs is an individual animal. Statistical significance was taken as ∗p < 0.05, ∗∗p < 0.01, ∗∗∗p < 0.001, and ∗∗∗∗p < 0.0001, as determined by unpaired two-tailed t-test, one-way repeated measures ANOVA, two-way repeated measures ANOVA, three-way repeated measures ANOVA or mixed-effect analysis followed by Tukey’s post hoc tests as appropriate. Statistical analyses were performed in GraphPad Prism 9.0 (Graphpad, La Jolla, CA).

## Results

To adapt the punishment risk task from rats to mice, we first sought to replicate the task as closely as possibly to the rat protocol. However, to accommodate size differences between the species we first reduced the total number rewards to from 150 to 45 pellets. This change in reward number also required changes to the contingencies used for the probabilistic punishment for Block 2 (0.06 to 0.066) and Block 3 (0.10 to 0.133). We initially kept the shock amplitude (0.3 mA) and duration (300 ms) the same[11]. To test if these changes would be sufficient to port the behavior to mice, we transferred animals from our breeding facility to our laboratory facility and allowed them to acclimate for one week. We next food restricted the animals to 90% of their initial body weight to increase motivation to seek food rewards during the light cycle (**Figure 1A)**. Next, animals were trained to self-administer sweetened food pellets on a FR1 schedule for 10 days (or until animals collected 45 pellets on 2 consecutive days) (**Figure 1B, Figures E-I)**. Following FR1 training with no punishment contingency, animals were tested with the adapted probabilistic punishment schemes (**Figure 1C**). Briefly, the punishment risk task (PRT) as adapted for mice requires mice learning to nosepoke in response to a cue to receive a sucrose pellet under a FR1 schedule of reinforcement for a total of 45 trials. During PRT this conditioning behavior is separated into 3 blocks (15 trials each), with each having increasing probability of nosepoke-contingent foot shock (0, 0.066, and 0.133) (**Figure 1C**). This design yields two discrete outcomes: 1) nosepoke latency, which can be considered the primary independent variable of PRT, quantifies the delay in instrumental action following cue onset; and 2) reward latency, a secondary independent PRT variable that quantifies the time an animal takes to retrieve the reward after it is delivered (**Figure 1C**). On this first test session animals increased latency to respond to the cue (**Figure 1D**, left). However, four out of the fourteen animals we tested did not complete all 45 trials and were excluded the analysis (**Figure 1J**). To determine whether this increased nosepoke response latency would carry over to future tests, we tested the animals again three days later. Encouragingly, on Test Day 2, there was no retained increased latency in *Block 1* and the increased latency in *Block 3* was recapitulated following the probabilistic shock exposure (**Figure 1D**, right). Unfortunately, however, far fewer mice completed the second test day (n=5/14; **Figure 1J)**. Considering almost all the animals completed all 45 trials when there was no probabilistic punishment, we presumed the shock amplitude was likely too great for the assay to be effective and repeatable in mice at 0.3 mA.

**Figure 1:**
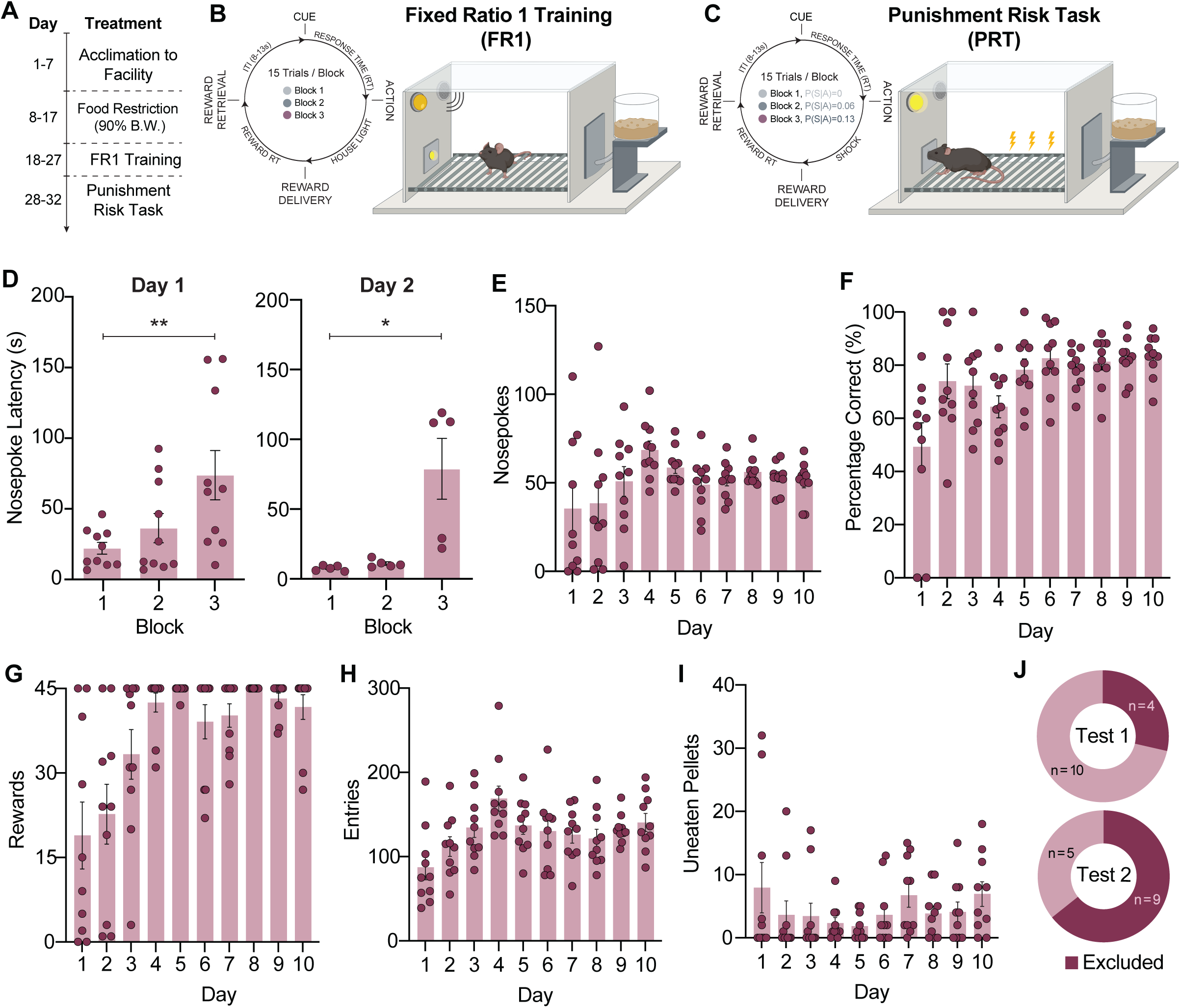
Initial study testing the Punishment Risk Task in male and female mice. **(A)** Calendar of behavioral testing **(B&C)** Diagrams depicting specifics of behavioral tasks during training (B) and test days (C). **(D)** Average nosepoke latency increases as probability of risk increases and is a repeatable behavioral test (Day 1: main effect of block, ** *p<0.01, n=10; Day 2: main effect of block, *p<0.05, n=5. **(E)** Average nosepokes during training days (1-10). **(F)** Percentage of correct nosepokes (Rewarded nosepoke / total nosepokes) during training days (1-10). **(G)** Average number of rewards received during training days (1-10). **(H)** Average number of pellet hopper entries during training days (1-10). **(I)** Average number of uneaten pellets left over during training days (1-10). **(J)** Percentage of animals excluded from analysis due to incomplete PRT test (Top: Test Day 1, Bottom: Test Day 2). Data are presented as mean ± SEM with dots reflecting individual subjects. For comparisons and exact p-values see **Table S1.**

To directly test the effect of shock amplitude on PRT performance, we used a second cohort of naïve animals. We randomly assigned each mouse to be singly-housed or kept in their group housing given future use of PRT would likely employ neurophysiological approaches that may require single housing (**Figure 2A**). To determine whether this adult social isolation alters baselined approach-avoidance behaviors, animals were tested in canonical exploration-based assays prior to PRT testing (**Figure S1**). In the open field test (OFT), there were no differences in exploration of the center zone between group-and single-housed animals or between sexes within or between these housing conditions **(Figure S1A-C)**. However, single-housed mice traveled significantly greater average distance than group-housed mice. Specifically, single-housed female mice moved significantly greater distance during testing compared to group-housed female and male mice **(Figure S1D)**. Though trending, these isolated mice were not significantly different from single-housed male mice. Further, there was no difference in the time spent in the open arms for group-and single-housed animals in the elevated plus maze (EPM) **(Figure S1E-F)**. Additionally, we did not observe any differences in open arm exploration between sexes or these housing conditions **(Figure S1G)**. Again, however, single-housed mice traveled greater average distance than group-housed mice, but sex did not affect distance traveled in the EPM **(Figure S1H).** Although social isolation (i.e., single housing) can impact approach-avoidance behaviors, we show here that this did not occur when the isolation is performed in adulthood as described previously[20].

**Figure 2:**
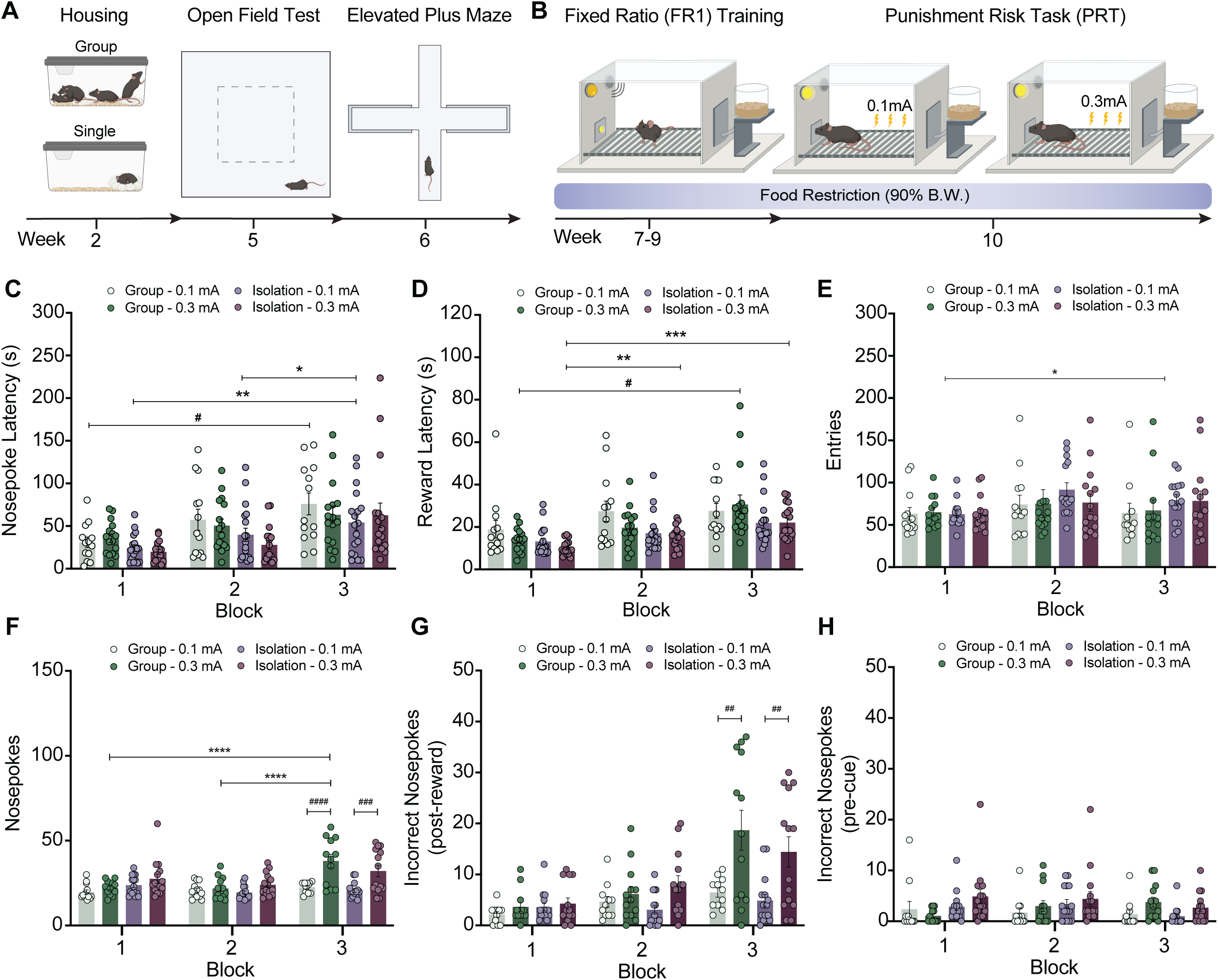
Punishment risk increases caution during goal-directed actions with differential effects dependent on shock amplitude and housing conditions. **(A)** Cartoon depicting the calendar of social isolation and canonical behavioral testing. **(B)** Calendar for Punishment Risk Task (PRT) training and testing. **(C)** Increasing probability of punishment increases average latency to nosepoke following cue presentation in both group-housed and single-housed animals regardless of shock intensity (0.1 or 0.3 mA) with post hoc comparisons (Group: 0.1mA-Block 1 vs 0.1mA-Block 3, # *p* < 0.05; Isolation: 0.1mA-Block 1 vs 0.1mA-Block 3, ** *p* < 0.01; Isolation: 0.1mA-Block 2 vs 0.1mA-Block 3, * *p* < 0.05). **(D)** Increasing probability of punishment increases average latency to retrieve pellet in both group-housed and single-housed animals regardless of shock intensity, post hoc comparisons (Group: 0.3mA-Block 1 vs 0.3mA-Block 3, # *p* < 0.05; Isolation: 0.3mA-Block 1 vs 0.3mA-Block 2, ** *p* < 0.01; Isolation: 0.3mA-Block 1 vs 0.3mA-Block 3, *** *p* < 0.001). **(E)** Increasing probability of punishment increases average number of reward hopper entries in all animals (*p* < 0.05) **(F)** Greater intensity shock (0.3 mA) increases the average number of total nosepokes in Block 3 for both group-housed animals (Group-Block 1 vs. Group-Block 3, **** *p* < 0.0001; Group-Block 2 vs. Group-Block 3, **** *p* < 0.0001; Group (0.1 mA)-Block 3 vs. Group (0.3 mA)-Block 3, #### *p* < 0.0001; and single-housed animals (Isolation (0.1 mA)-Block 3 vs. Isolation (0.3 mA)-Block 3, ### *p* < 0.001). **(G)** Greater intensity shock (0.3 mA) increases the average number of incorrect nosepokes following reward presentation (Group (0.1 mA)-Block 3 vs. Group (0.3 mA)-Block 3, ## *p* < 0.01 and single-housed animals (Isolation (0.1 mA)-Block 3 vs. Isolation (0.3 mA)-Block 3, ## *p* < 0.01) **(H)** Housing conditions and shock intensity do not have any effect on incorrect nosepokes during the intertrial period prior to cue presentation. Data are presented as mean ± SEM with dots reflecting individual subjects. Data are presented as mean ± SEM with dots reflecting individual subjects. For comparisons and exact p-values see **Table S1.**

Following OFT and EPM testing, animals were then food restricted to 90% of their initial body weight and trained to self-administer sweetened food pellets for PRT testing (**Figure 2B**). This involved training on a FR1 schedule for 10 days (or until animals collected 45 pellets on 2 consecutive days) (**Figure S2B-D, H,I)**. Following FR1 training with no punishment contingency, animals were assigned to specific shock amplitudes in a counterbalanced manner and tested in PRT with the adapted probabilistic punishment schemes (**Figure 2B**). Each test required the animals complete all three blocks to be included in behavioral data analysis **(Figure S2A**). Prior to shock introduction, animals tested with no punishment probability had similar nosepoke latency across blocks and between groups (**Figure S2E).** Despite no difference in the motivation to respond to the cue, these groups were shown have an increase in reward latency as they progressed through the test (**Figure S2F**) and did not consume all pellets received (**Figure S2G**). This likely indicates some effect of time or satiety on the motivation to retrieve a reward following successful operant conditioning behavior without a probabilistic punishment.

To establish an effective punishment without producing fear-conditioned behavior, we tested two different 300 ms shock amplitudes (0.1 mA and 0.3 mA). We observed a significant effect of block and housing on nosepoke latency, but no significant effect of shock amplitude (**Figure 2C**). This indicates that the animals were slower to respond as the probability of punishment increases in each block, but there was no significant interaction between any of the factors. Closer analysis revealed significant differences between average nosepoke latency in *Block 1* and *Block 3* for 0.1 mA shock in both isolated animals and grouped animals. Additionally, average nosepoke latency was significantly increased from *Block 2* to *Block 3* for 0.1 mA shock in isolated animals, but not group-housed animals (**Figure 2C**). We did not observe any significant differences comparing blocks between isolated and grouped animals for 0.3 mA shock. Further, we observed a significant effect of block and housing on reward latency, but no significant effect of shock amplitude (**Figure 2D**). This indicates that animals took more time to retrieve the reward as the probability of punishment increases in each block. There was, however, no significant interaction between any of the factors. There were significant differences between average reward retrieval latency in *Block 1* and *Block 3* for 0.3 mA shock for both isolated and grouped animals. Specifically, average reward retrieval latency was increased between *Block 1* and *Block 2* for 0.3 mA shock in isolated animals, but not group-housed animals (**Figure 2D**). We did not observe any relevant significant differences between isolated and grouped housed animals for 0.3 mA shock or any significant differences in any block or housing condition for 0.1mA. Together, these results suggest that higher intensity shock may be more likely to increase reward retrieval latency alongside latency to respond to cue presentation. As previously described[11], this increased reward retrieval latency confounds clear interpretation of the hesitancy to nosepoke for reward following the cue. Additionally, we observed significant increase in reward retrieval entries between *Block 1* and *Block 3*, yet we did not observe any significant effects of housing condition or shock amplitude **(Figure 2E).** Further, our data demonstrated that higher intensity shock drives increased nose-poking behavior during *Block 3* for group and single-housed animals (**Figure 2F**). Greater intensity shock (0.3 mA) increases the average number of total nosepokes in *Block 3* for both group animals and single-housed animals **(Figure 2F**). Further analysis revealed that animals committed significantly more incorrect nosepokes during the period following the reward presentation in *Block 3* when receiving 0.3mA shock **(Figure 2G)** while having no significant difference in incorrect nosepokes during the intertrial period prior to the cue presentation **(Figure 2H)**. Given this propensity of animals to have a stronger behavioral response to 0.3mA footshock, we also examined nosepoke latency for the trial immediately following the shock presentation. Here we did not observe any significant change in nosepoke latency for trials immediately following shock nor any significant differences due to housing or shock intensity **(Figure S3A)**. To minimize any influence of shock intensity on behavioral responding, further examination of PRT with drug treatments only used a 0.1 mA shock amplitude.

Following Park and Moghaddam [11], we then used the benzodiazepine anxiolytic, diazepam, to test whether the increased nosepoke latency in *Block 3* can be reversed similar to anxiety states observed in humans **(Figure 3A)**. Here, diazepam treatment (2 mg/kg) significantly blunted the increase in nosepoke latency in both single-and group-housed mice compared to within-subject saline treatment (**Figure 3B-C**). We observed similar results for the average reward retrieval latency in both group-and single-housed mice. Reward retrieval latency increased as punishment risk increased while diazepam treatment prevented this increase in both groups (**Figure 3D-E**.) These data suggest that anxiolytic diazepam prevents the increased latency to respond to cue presentations during sessions with higher punishment probability. These experiments also demonstrated that saline injection alone influences reward retrieval latency which was not observed in previous 0.1 mA shock PRT testing. Despite repeated handing prior to drug treatment experiments, this may be due to added stress that injection procedures may have on animals that were not used in previous testing. Further behavioral analysis revealed diazepam treatment to differentially affect the average number of nosepokes per block for single-housed animals (**Figure 3F**). Specifically, diazepam treatment increases average nosepokes for single-housed animals in *Block 1* as compared to saline treatment and as compared to group-housed animals. Single-housed animals have greater average nosepokes in *Block 1* following saline and diazepam treatment (**Figure 3F**). Further analysis revealed that single-housed animals committed significantly more incorrect nosepokes during the intertrial period prior to the cue in *Block 1* following diazepam treatment **(Figure 3H)**, while having no significant difference in incorrect nosepokes following the reward presentation **(Figure 3G)**. We also examined nosepoke latency for the trial immediately following the shock presentation. Here we did not observe any significant change in nosepoke latency for trials immediately following shock nor any significant differences due to housing or diazepam treatment **(Figure S3B)**.

**Figure 3:**
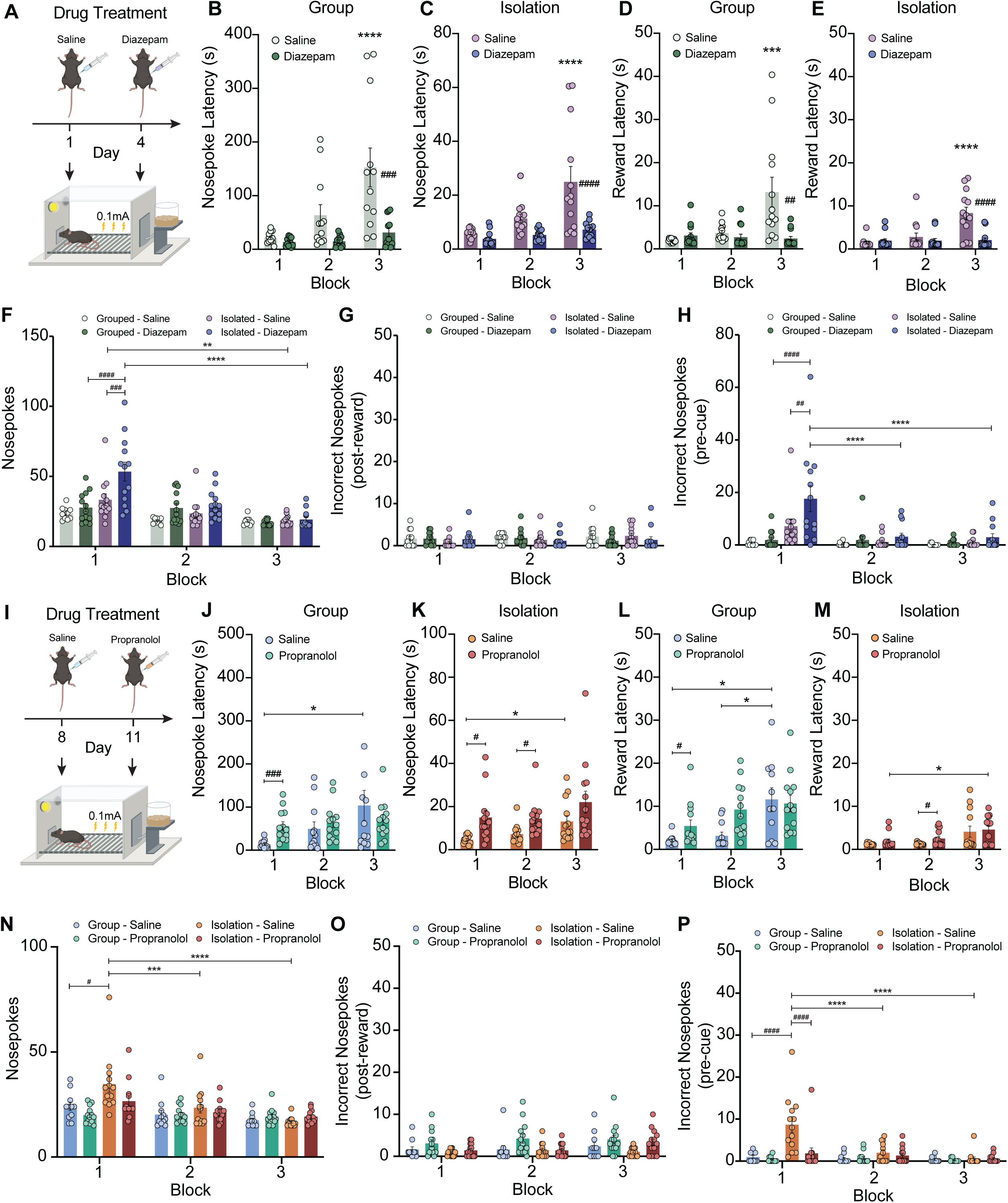
Diazepam and propranolol prevent punishment risk-associated response latency increase during goal-directed actions. **(A)** Calendar for diazepam treatment and Punishment Risk Task (PRT) testing. **(B)** Diazepam (2 mg/kg) treatment reduces average latency to nosepoke among group-housed animals as compared to saline treatment; Saline-Block 1 vs Saline-Block 3, **** *p* < 0.0001; Saline-Block 3 vs. Diazepam-Block 3, #### *p* < 0.0001 **(C)** Diazepam treatment reduces average latency to nosepoke among single-housed animals as compared to saline treatment; Saline-Block 1 vs Saline-Block 3, **** *p* < 0.0001; Saline-Block 3 vs. Diazepam-Block 3, #### *p* < 0.001. Note the change in scale on y-axes between **B** & **C. (D)** Diazepam treatment reduces average latency to retrieve reward pellet among group-housed animals as compared to saline treatment; Saline-Block 1 vs Saline-Block 3, *** *p* < 0.001; Block 3-Saline vs. Diazepam-Block 3, ## *p* < 0.01. **(E)** Diazepam treatment reduces average latency to retrieve reward pellets among single-housed animals as compared to saline treatment; Saline-Block 1 vs Saline-Block 3, ** *p* < 0.01; Saline-Block 3 vs. Diazepam-Block 3, # *p* < 0.05. **(F)** Diazepam treatment in single-housed animals increases the average number of nosepokes completed in the first block, prior to any probabilistic punishment, as compared to saline treatment; Saline-Block 1 vs Diazepam-Block 1, ###, *p* < 0.001. This effect is also observed when compared to diazepam-treated group-housed animals; Group, Diazepam – Block 1 vs. Isolation, Diazepam-Block 1, #### *p* < 0.0001. Diazepam-treated single-housed animals also perform more nosepokes in Block 1 compared to Block 3; Diazepam-Block 1 vs. Diazepam-Block 3, **** *p* < 0.0001. Saline-treated single-housed animals also perform more nosepokes in Block 1 compared to Block 3; Saline-Block 1 vs. Saline-Block 3, ** *p* < 0.01. Diazepam treatment has no effect on group-housed animals; Saline-Block 1 vs. Diazepam – Block 1, *ns, p* = 0.99; Saline-Block 2 vs. Diazepam – Block 2, *ns, p* = 0.70; Saline-Block 3 vs. Diazepam – Block 3, *ns, p* = 0.99. **(G)** Housing conditions and drug treatment have no effect on incorrect nosepokes performed during the period following the reward presentation. **(H)** Diazepam treatment in single-housed animals increases the average number of incorrect nosepokes completed during the inter-trial period before cue presentation in the first block, prior to any probabilistic punishment, as compared to saline treatment; Saline-Block 1 vs Diazepam-Block 1, ##, *p* < 0.01. This effect is also observed when compared to diazepam-treated group-housed animals; Group, Diazepam – Block 1 vs. Isolation, Diazepam-Block 1, #### *p* < 0.0001. Diazepam-treated single-housed animals also perform more incorrect nosepokes in Block 1 compared to Block 2 and Block 3; Diazepam-Block 1 vs. Diazepam-Block 2, **** *p* < 0.0001, Diazepam-Block 1 vs. Diazepam-Block 3, **** *p* < 0.0001. Diazepam treatment has no effect on group-housed animals. **(I)** Calendar for propranolol treatment and Punishment Risk Task (PRT) testing. **(J)** Propranolol (10 mg/kg) treatment prevents increased average latency to nosepoke among group-housed animals as compared to saline treatment; Saline-Block 1 vs Saline-Block 3, * *p* < 0.05; Propranolol-Block 1 vs. Propranolol-Block 3, *ns, p* = 0.92. Propranolol treatment increases nosepoke latency during Block 1; Saline-Block 1 vs. Propranolol-Block 1, ### *p* < 0.001. Note the change in scale on y-axes between **J** and **K. (K)** Propranolol (10 mg/kg) treatment prevents increased average latency to nosepoke in single-housed animals as compared to saline treatment; Saline-Block 1 vs. Saline-Block 3, ** *p* < 0.01; Propranolol-Block 1 vs Propranolol-Block 3, ns, *p* = 0.99. **(L)** Propranolol (10 mg/kg) treatment has no effect on average latency to retrieve reward pellet among group-housed animals as compared to saline treatment; Saline-Block 1 vs Saline-Block 3, *** *p* < 0.001. Propranolol treatment increases reward retrieval latency during Block 2; Saline-Block 2 vs. Propranolol-Block 2, # *p* < 0.05. **(M)** Probabilistic punishment increases latency to retrieve reward pellet in single-housed animals; Saline-Block 1 vs. Saline-Block 3, * *p* < 0.05; Propranolol-Block 1 vs Propranolol-Block 3, #, *p* < 0.05. **(N)** Saline treatment in single-housed animals increases the average number of nosepokes completed in the first block, prior to any probabilistic punishment, as compared to group-housed animals; Group, Saline-Block 1 vs Isolation, Saline-Block 1, #, *p* < 0.05. This effect is also observed when compared to blocks 2 and 3 in saline-treated, single-housed animals; Saline-Block 1 vs. Saline-Block 2, *** *p* < 0.001; Saline-Block 1 vs. Saline-Block 3, **** *p* < 0.0001. **(O)** Housing conditions and drug treatment have no effect on incorrect nosepokes performed during the period following the reward presentation in propranolol-treated animals. **(P)** Single-housed animals have greater number of incorrect nosepokes completed during the inter-trial period before cue presentation in the first block as compared to group-housed animals; Group, Saline-Block 1 vs. Isolation, Saline-Block 1, #### *p* < 0.0001. Propranolol treatment in single-housed animals prevents this increase in incorrect nosepokes completed prior to cue presentation in the first block, as compared to saline treatment; Saline-Block 1 vs Propranolol-Block 1, ####, *p* < 0.0001. This effect in single-housed animals is only observed in the first block of the task; Saline-Block 1 vs Saline-Block 2, **** *p* < 0.0001, Saline-Block 1 vs Saline-Block 3, **** *p* < 0.0001. Propranolol treatment has no effect on group-housed animals. Data are presented as mean ± SEM with dots reflecting individual subjects. Experiments used 0.1mA shock for probabilistic punishment. For comparisons and exact p-values see **Table S1.**

To determine whether another class of anxiolytic could reverse PRT, we next tested propranolol, a non-selective β-adrenergic receptor antagonist with anxiolytic properties in humans and mice **(Figure 3I)**[17,18,21,22]. Here, increased nosepoke latency across blocks was prevented following propranolol (10 mg/kg) treatment (*Block 1 vs Block 3*) in group housed animals (**Figure 3J**). However, propranolol treatment increased nosepoke latency in *Block 1.* Similarly, in singly-housed mice, we observed a significant in nosepoke latency between mice across blocks in saline-treated animals, yet propranolol treatment increased nosepoke latency in *Block 1 and Block 2* compared to saline treatment (**Figure 3K**). Again, mice have increased average nosepoke latency in *Block 3* compared to *Block 1* following saline treatment, but these latencies were not significantly different in propranolol-treated animals. These effects were also observed in animals’ average latency to retrieve the reward. Group and single-housed animals had statistically significant increases in reward retrieval latency following saline treatment, with propranolol treatment reducing the average reward retrieval latency in *Block 3* as compared to saline treatment for isolated mice, but not group-housed mice **(Figure 3L-M)**. Single-housed animals have greater average nosepokes in *Block 1* following saline treatment while propranolol treatment had no significant effect on committed nosepokes (**Figure 3N**). Further analysis revealed that saline treated, single-housed animals committed significantly more incorrect nosepokes during the intertrial period prior to the cue in *Block 1* following diazepam treatment, while having no significant difference in incorrect nosepokes following the reward presentation **(Figure 3O&P)**. Finally, we examined nosepoke latency for the trial immediately following shock presentations in group-and single-housed animals. Here we observed an effect of housing on nosepoke latency for trials immediately following shock, regardless of the number of shock presentations. Multiple comparisons revealed that in propranolol-treated, group-housed animals have greater average nosepoke latency following the third shock presentation as compared to propranolol-treated, single-housed animals differences due to housing or diazepam treatment **(Figure S3C)**. These data suggest a unique role for β-adrenergic receptor signaling during PRT that prevent appropriate integration of action-punishment contingencies.

To determine whether PRT testing alters locomotor activity across blocks as a proxy for shock-induced freezing, we used video recording and analysis (**Figure 4A)**. Mice have greater movement velocity during *Block 1* as compared to *Block 2* and *Block 3*, but no significant difference in immobility time (**Figure 4B-C**). Further, mice spend a greater percentage of time near the food hopper compared to the nosepoke port in all blocks regardless of shock probability (**Figure 4D**). Lastly, mice show a reduction in average number of entries (per minute) into the zones near the hopper and the nosepoke port over time. (**Figure 4E**).

**Figure 4:**
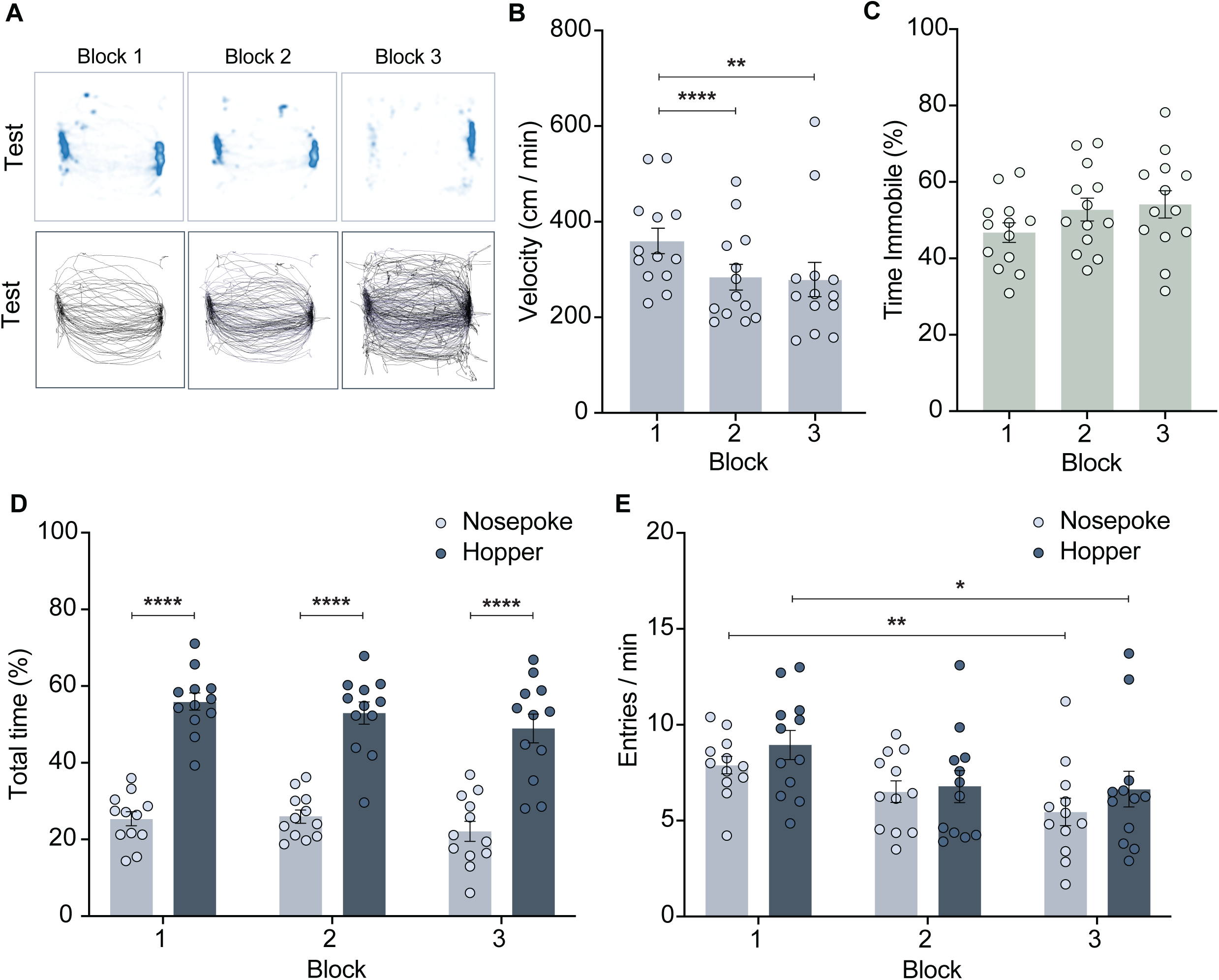
Punishment risk and task related activity behaviors. **(A)** Representative heatmaps and traces for each block during PRT. **(B)** Average velocity reduces throughout test session (Block 1 vs Block 2, **** *p* < 0.0001; Block 1 vs Block 3, **** *p* < 0.01). **(C)** Punishment risk does not affect time spent immobile in single-housed animals (ns). **(D)** Regardless of block, animals spend a greater percentage of session in hopper zone compared to nosepoke zone (Block 1-Nosepoke vs Block 1-Hopper, **** *p* < 0.0001; (Block 2-Nosepoke vs Block 2-Hopper, **** *p* < 0.0001; Block 3-Nosepoke vs Block 3-Hopper, **** *p* < 0.0001). **(E)** Average zone entries per minute reduces throughout test session (Block 1 Nosepoke vs Block 3 Nosepoke, **** *p* < 0.0001; Block 1-Hopper vs Block 3-Hopper, **** *p* < 0.01). Data are presented as mean ± SEM with dots reflecting individual subjects. Experiments used 0.1mA shock for probabilistic punishment. For comparisons and exact p-values see **Table S1.**

Finally, to better understand the construct and content validity of PRT, we continued to test animals under a typical control treatment condition (saline injection, i.p.). Despite this injection having a small impact on stress responsivity, this procedure does not impact or influence the effect of PRT testing (**Figure 5A-B; see also Figure 3**). Probabilistic punishment significantly increased the trial-to-trial variability in nosepoke latency for single-housed mice, yet this variability was not significant in group-housed mice **(Figure 5D-E)**. This may be due to group-housed animals having larger variance within all blocks, generally **(Figure 5C)**. In contrast, the mean nosepoke latency and the variance nosepoke latency did not differ across blocks when there is no probability of shock across blocks **(Figure 5F)**. To determine if mice learn the nosepoke-shock contingency or, instead, increase nosepoke latency as a general consequence of unpredictable shock, we compared nosepoke latencies prior to the first shock delivery in *Block 2* and *3*. Isolated mice increased nosepoke latency in *Block 2* and *3,* however, this increase was not significant for group-housed mice. In contrast, this increase was significant in *Block 3* before the first shock in single-housed mice **(Figure 5G)** suggesting that single housing could influence learning of action-punishment contingencies from minimal previous experience.

**Figure 5:**
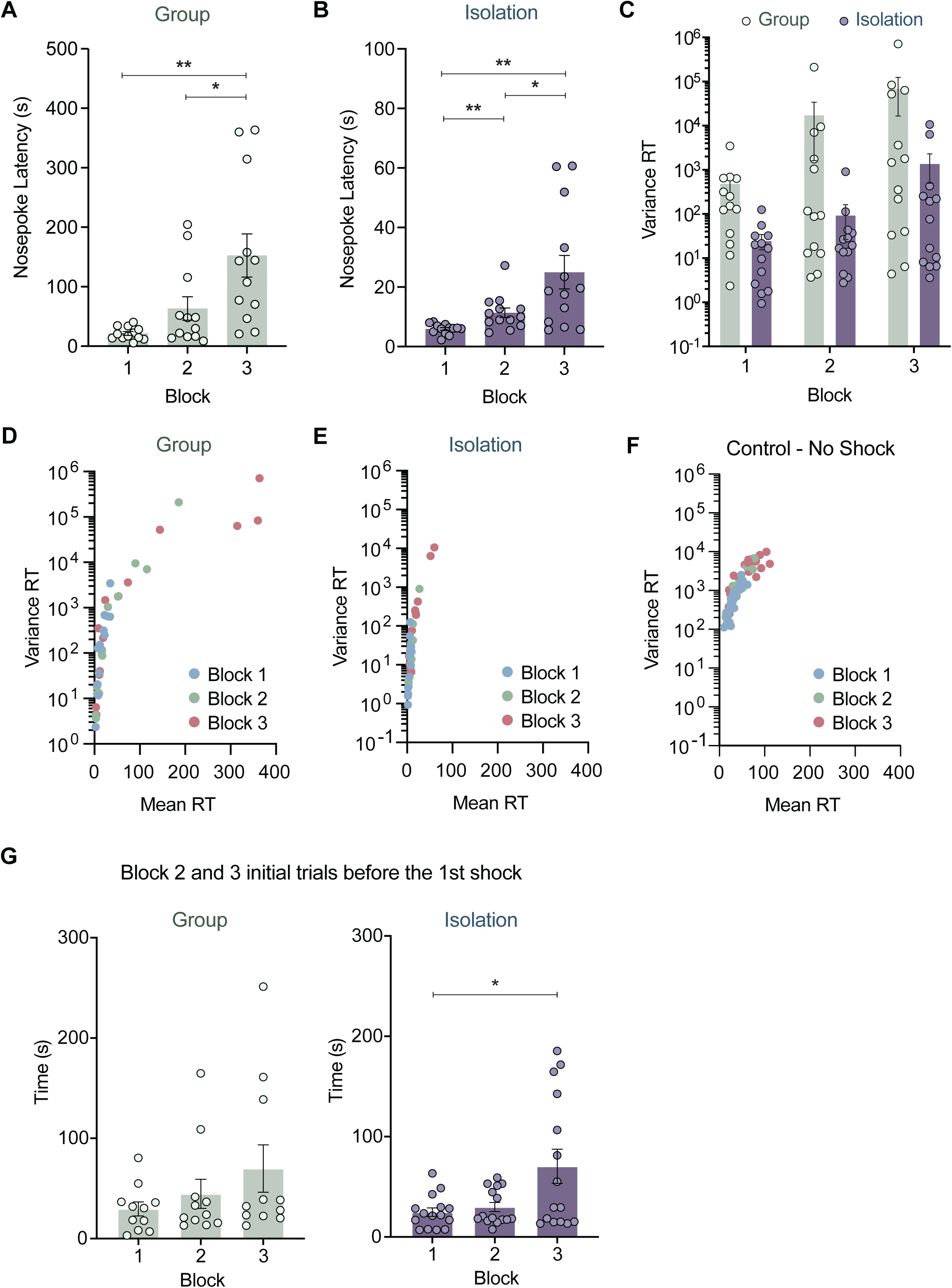
Punishment risk increases variance for trial-to-trial nosepoke latency. **(A)** Increased average nosepoke latency with increasing punishment risk following saline treatment among group-housed animals (Block 1 vs Block 3, ** *p* < 0.01; Block 2 vs Block 3, * *p* < 0.05) and **(B)** single-housed animals (Block 1 vs Block 3, ** *p* < 0.01; Block 1 vs Block 2, ** *p* < 0.01; Block 2 vs Block 3, * *p* < 0.05). Note change in scale for y-axes in **A** and **B**. **(C)** Average trial-to-trial variance for each block in group and single-housed animals. **(D)** Colored dots represent individual animals’ variance in nosepoke latency across trials in different blocks as a function of mean nosepoke latency for group and single-housed animals. **(E)** Individual mean and variance in nosepoke latency of all animals across trials (combined group and isolation animals) and **(F)** no-shock control session for all animals. **(G)** To determine whether mice learn the low-probability nosepoke-shock contingency, we compared the nosepoke latency of the pre-shock trials in Block 2 and Block 3 for group and single-housed animals (* p<0.05). Data are presented as mean ± SEM with dots reflecting individual subjects. For comparisons and exact p-values see **Table S1.** For **D-F**, data are presented as corresponding points individual subjects’ variance and mean for each block.

## Discussion

We have translated an important rat behavioral task into mice and demonstrated its utility as a murine model for assessing repeatable anxiolytic-sensitive hesitancy behavior. Importantly, this model’s task-oriented design allows the interrogation of behavioral reinforcement and passive avoidance, both important behavioral measures to more accurately quantify murine behavioral phenotypes related to negative affective disorders such as Generalized Anxiety Disorder. Critically, this test can be repeated multiple times under multiple conditions enabling long-term studies, increasing within-subject power, and reducing the overall number of animals needed to test. In addition to porting this assay to both sexes of mice, our study design included evaluating the effects of chronic isolation in adulthood. This approach allowed us to further determine whether this model would detect any anxiety-or motivation-related effects of adult single housing, a condition sometimes necessitated in behavioral and circuit neuroscience. Furthermore, underlying neural circuit dynamics can be probed at neurophysiological timescales with this approach[11].

Although mice successfully completed the task following modifications, many mice did not consume all dispensed pellets (**Figure S2G**). Alas, this number of rewards (15 per block) was the minimum possible, given that the probability of shock needed for *Block 2* resulted in only one shock. To avoid this restriction, future studies could use liquid rewards (e.g., sucrose solutions) and with a limited-access lickometer. This could enable more flexibility to change probabilities and number of trials per block, helping minimize any satiety or motivational constraints and maintain responding and consumption throughout the entirety of the task. However, the explicit goal of this study was to maintain validity in translating this task to mice with as little deviance from the original rat model as possible. Further, we tested an additional shock intensity, 0.1 mA, along with the previously demonstrated 0.3 mA shock. Here we found that, as with previous rat studies[11,23,24], mice only exhibit higher latency to act on the operant nosepoke port following cue presentations under conditions of probabilistic punishment. This effect was most prominent for sessions using 0.1 mA shock in both group-and single-housed animals **(Figure 2C)**. However, despite not having any significant difference in nosepoke latency for group-and single-housed mice, 0.3 mA shock blocks had a significant effect on reward retrieval latency in both group and single-housed mice in *Block 3* **(Figure 2D)** and increased the likelihood of animals not completing the task (**Figure 1J**). Given this striking discrepancy in behavioral outcome, we selected 0.1 mA shock intensity for subsequent anxiolytic treatments (**Figure 3 & Figure 4**).

Importantly, as previously demonstrated in rats[11], diazepam treatment abolishes the behavioral response to increased punishment probability. Previous studies have demonstrated diazepam as an anxiolytic in C57BL/6 mice[25]. However, a recent study has suggested that these effects are largely driven by sedation in open field and elevated plus maze testing[26]. Despite this conflicting evidence, our data do not suggest any sedative effect given the animals’ propensity to continue to perform the task and, importantly, do so even more efficiently. For both group-and single-housed animals, we observed an increase in nosepoke latency and reward latency with saline treatment, while diazepam treatment reduced these changes in response latency (**Figure 3B-E**). These data suggest that increased latency in responding is indicative of an anxiogenic state given that diazepam only reduces nosepoke latency during probabilistic punishment in *Block 3*. Additionally, it is unlikely that diazepam affects the sensitivity to footshock as previous research has demonstrated neither sex nor diazepam treatment to have any effect on foot shock sensitivity[27]. Importantly, we did not observe any effect of animal housing on the canonical anxiety-related behavioral assessments of exploratory approach-avoidance conflict (**Figure S1A,B,E,F**), despite previous work reporting reduced anxiety-like behavior in female rodents[20,28]. However, in our hands, social isolation increased distance traveled in both OFT and EPM testing, contrary to other results[20]. Despite these conflicting data, we propose that the effects observed in our experiments are not at odds with previous study, but inform potential differences in implementing this assay in different species[5,12].

Further, isolated mice had lower nosepoke latencies, generally (see **Figure 5A-B**), with less variance in responding during probabilistic punishment. As with rats[11], changes in nosepoke latency during probabilistic punishment blocks parallels changes in the variance of response latencies within those blocks in mice (i.e. as the probability of punishment increases, the mean and variance for nosepoke latency for the block increases; **Figure 5C-E**), where responding during *Block 3* shows the greatest variance among the highest mean responding. This increased variance could be characterized as a measure of “hesitancy” or “caution” during this reward seeking behavior. Indeed, a recent study found that mice delayed their action for a signaled reward when a new, unsignaled rule that punished the action was added[30]. However, the interpretation from that study suggested that this “caution” is not due to an anxiogenic state as it is unaffected by diazepam administration[30]. This discrepancy is likely due to multiple different factors: apparatus (shuttle box vs. operant box), animal state (water-restricted vs. food-restricted), reward (water vs. sweetened pellet), and the probability of punishment (1.00 vs 0.13). Our model examined this change in signal-response latency during a period of low probability of unsignaled punishment, while the other model relied on either explicit unsignaled action-contingent punishment or punishment without contingency. Given these differences, it is difficult to make direct comparisons of the “caution” demonstrated in these rodent behavioral models. One unique observation from our studies revealed that different strategies for reward seeking may develop as animals become familiar with the task. Initially, we observed increased nosepokes in both group-and single-housed animals during the third block of the 0.3mA testing that were attributed to “incorrect” nosepokes committed during the post-reward presentation period (**Figure 2F**). In later experiments, we observed this is “incorrect” nosepoke behavior only in single-housed animals during the first block, prior to any shock presentation (**Figure 3H, 3P**). Conversely, these incorrect nosepokes were performed during the inter-trial period prior to the cue presentation. This “impulsive” strategy where animals are eager to complete the task may be unique to single-house animals, but this behavioral phenotype has yet to be fully explored.

Recent work from the Moghaddam lab expanded on their previous findings to determine how diazepam affects this change in behavior[23]. Here they found that exposure to conflicting reward and punishment contingencies caused ventral tegmental area (VTA) and dorsomedial prefrontal cortex (dmPFC) neurons to have modified activity in the time period before the action and subsequent reward periods but did not change neuronal responses to punishment. Interestingly, treatment with diazepam potentiated VTA response to action and enhanced its correlated activity with the dmPFC, but still did not affect the neural response to the punishment. The authors propose that the encoding action-selective information, rather than punishment-selective information, drives the anxiogenic state observed in rats[23]. Overall, increased punishment risk likely disrupts goal-oriented neuronal activity, while diazepam administration enhances coordination of activity within this neurocircuitry to refocus this activity to influence and maintain goal-directed actions.

The main purpose of our study was to establish a model of anxiety-related behavior in mice that was analogous to the experience of risk uncertainty demonstrated in human populations [31–33]. Behavioral neuroscience assays that are used to study the neural basis of anxiety typically rely on exploratory behaviors in novel contexts[10,12]. Many of these tests can only be performed once per animal as the novelty of the environment is critical to the interpretation of the behavior. While these approaches have been incredibly valuable for understanding neurobiological mechanisms of acute fear and anxiety states[17,18,34–44], these assays do not explicitly model more complex situations where negative affective states can develop from learning that certain goal-directed actions have some risk of an aversive outcome. Alternatively, here we increase the probability of punishment while the animal performs an instrumental action to study this unique behavior. Humans report that unpredictable aversive stimuli, similar to what we show here in mice, generates increased anxiety that is disrupted by benzodiazepines[45–47]. Furthermore, similar approaches in rodents and non-human primates have shown benzodiazepine-mediated reversal is conserved across species, indicated that these approaches in animal models can provide powerful insight into the neurobiology of anxiety[11,23,24,48–50]. This translational impact is strengthened by recent human studies on the behavioral impact of punishment are driven by poor action-punishment contingency learning. Here we present a mouse assay to examine the learning of these contingencies that has significant potential for the study of anxiety mechanisms. Overall, these experiments demonstrate that effective implementation of the Punishment Risk Task (PRT) model is possible in mice and, with careful adaptation, mostly parallels the behavioral outcomes that have been seen in rat studies. We hope that this detailed study will enable wider adoption of this repeatable murine model for the preclinical study of anxiety-related behaviors.

## Supporting information

Med Associates Source Code

## Data availability

All data generated or analyzed during this study are included in the manuscript and the Source Data file.

## Code availability

The code used to run the Punishment Risk Task on Med Associates hardware is available in the Code file.

## Acknowledgements

We thank the other members of the Al-Hasani and McCall labs, particularly Marwa O. Mikati, Justin Woods, and Makenzie R. Norris, for helpful feedback on this project. This work was supported by the National Institutes of Health (R01NS117899, J.G.M.), the Brain & Behavior Research Foundation (NARSAD YI – 28565, J.G.M.), and the Rita Allen Foundation with help from the Open Philanthropy Project (J.G.M.). We would like to acknowledge biorender.com for figure cartoons and the Osage Nation, Missouria, Illinois Confederacy, and many other tribes as the ancestral, traditional, and contemporary custodians of the land where this work was conducted.

## Author contributions

K.E.P. and J.G.M conceived the project and designed the detailed experimental protocols. K.E.P. and S.C.H. performed the mouse experiments. K.E.P., J.S.A., S.C.H., C.H., and J.G.M performed the investigation and analyzed the data. K.E.P., and J.G.M wrote the paper. J.G.M acquired funding. J.G.M. provided research supervision and led overall project administration. All authors discussed the results and contributed to revision of the manuscript.

## Conflict of Interest

The authors declare no conflicts of interest.

---

**Figure S1:**
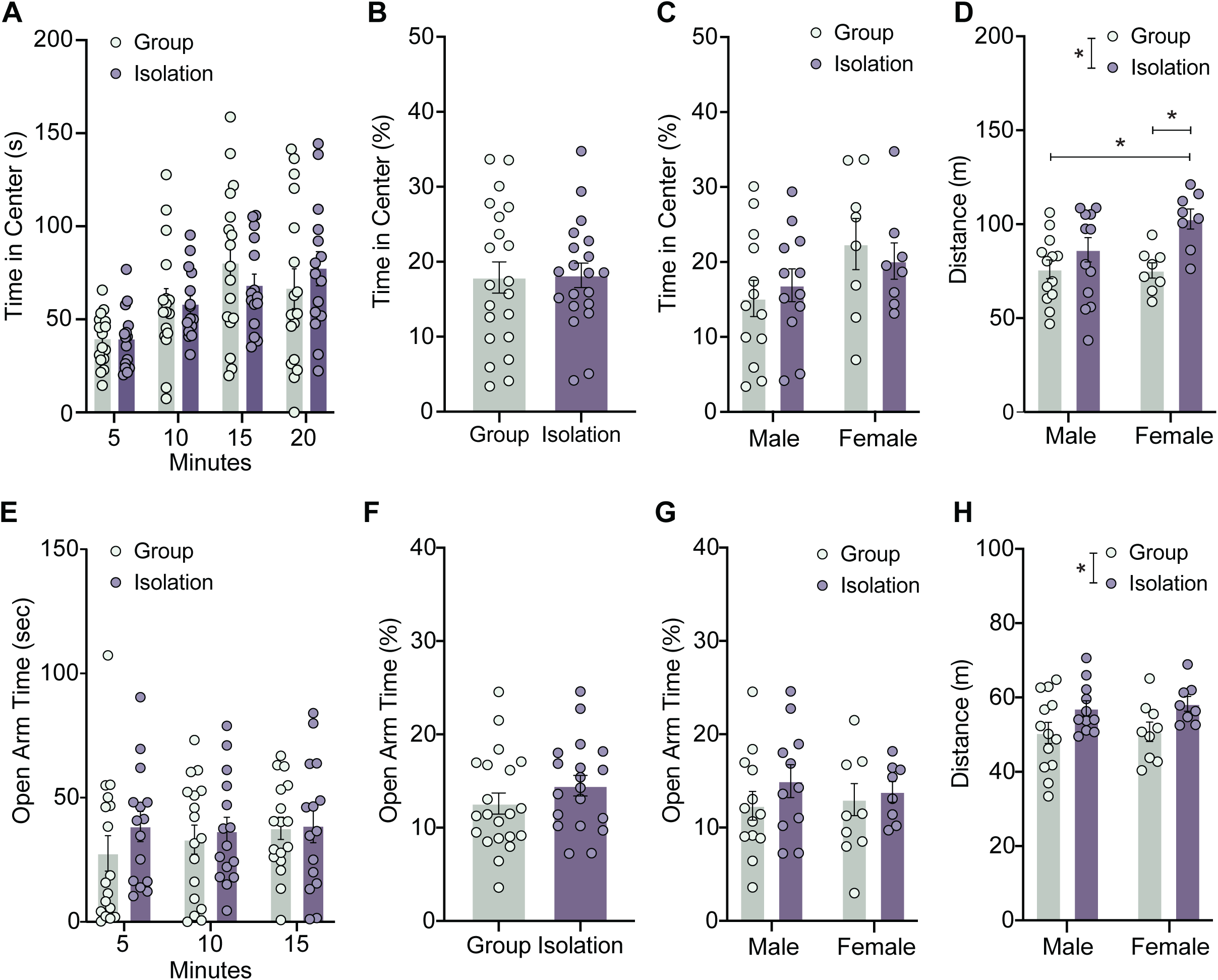
Adult social isolation has minimal effect on canonical exploratory behaviors. **(A)** Adult social isolation does not affect center zone exploration in open field test. **(B)** Male and female mice do not differ in the percentage of time spent exploring the center zone of open field test, regardless of housing. **(C)** Single-housed female mice have greater total distance traveled in the open field test compared to group-housed male mice (*p* < 0.05) and group-housed female mice (*p* < 0.05). **(D)** Heat maps depicting average time spent in open field test for group and single-housed mice. **(E)** Social isolation does not affect open arm exploration in elevated plus maze. **(F)** Male and female mice do not differ in the percentage of time spent exploring the open arms of elevated plus maze. **(G)** Isolation housing significantly increases total distance traveled in the elevated plus maze (Group vs Isolation, * *p* < 0.05), but male and female mice are not different within these groups. **(H)** Heat maps depicting average time spent in elevated plus maze for group and single-housed mice. Data are presented as mean ± SEM with dots reflecting individual subjects. For comparisons and exact p-values see **Table S1.**

**Figure S2:**
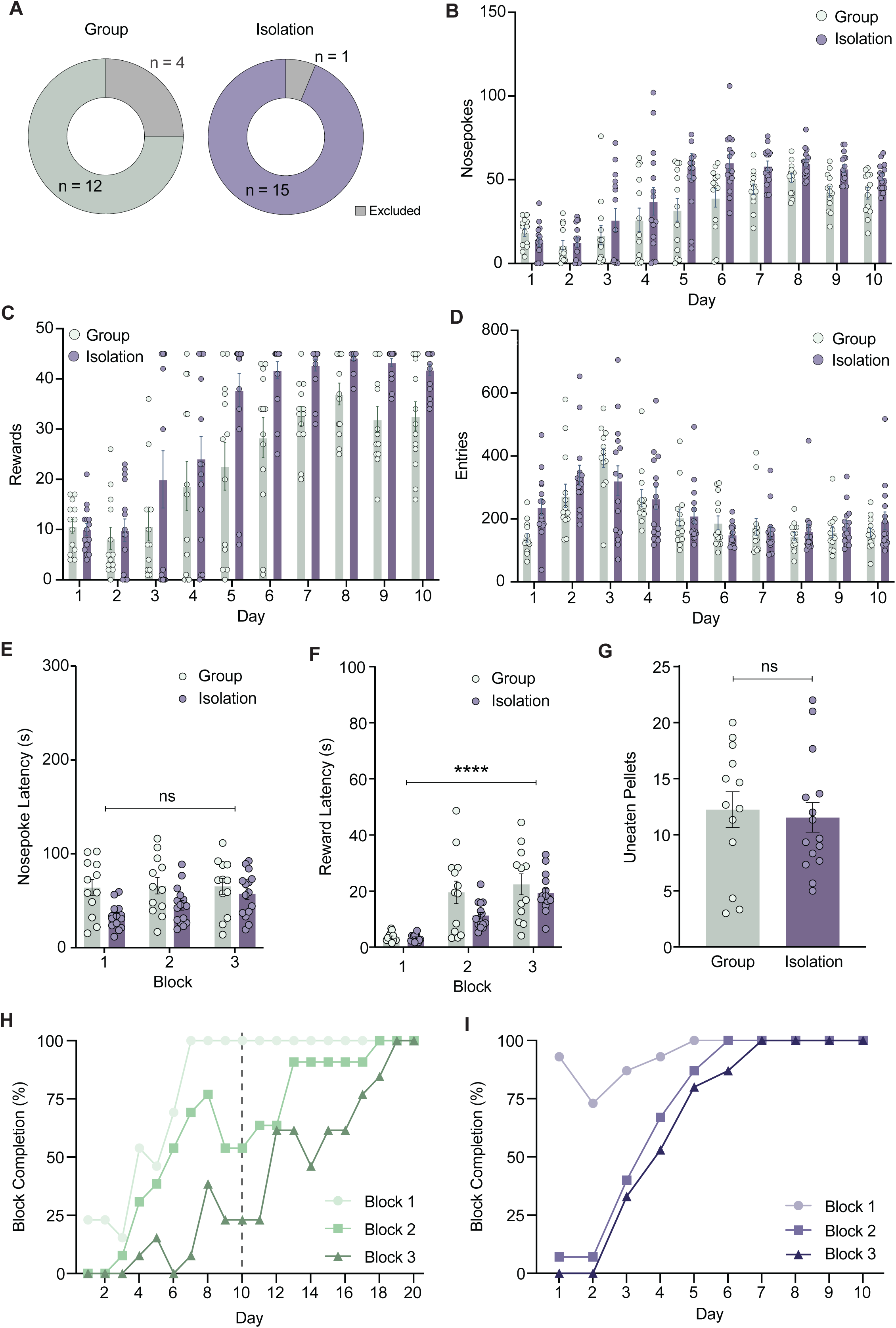
Punishment risk task training in group and single-housed mice. **(A)** Chart displaying total number of animals that did not successfully complete PRT training and were excluded from further analysis. **(B)** Average nosepokes during training days (1-10) in group and single-housed mice. **(C)** Average number of rewards received during training days (1-10). **(D)** Average number of pellet hopper entries during training days (1-10). **(E)** Animals do not display any significant change in average nosepoke latency during “no shock” test day. **(F)** Animals have increased average reward retrieval latency during no shock test day, Block 1 vs Block 3, **** *p* < 0.0001. **(G)** There is no significant difference in the average number of uneaten pellets left over during no shock test day (ns). **(H)** Percentage completion of blocks of rewards during training for group-housed mice. Dashed line indicates day single-housed animals completed training. **(I)** Percentage completion of blocks of rewards during training for single-housed mice. Data are presented as mean ± SEM with dots reflecting individual subjects. For comparisons and exact p-values see **Table S1.**

**Figure S3:**
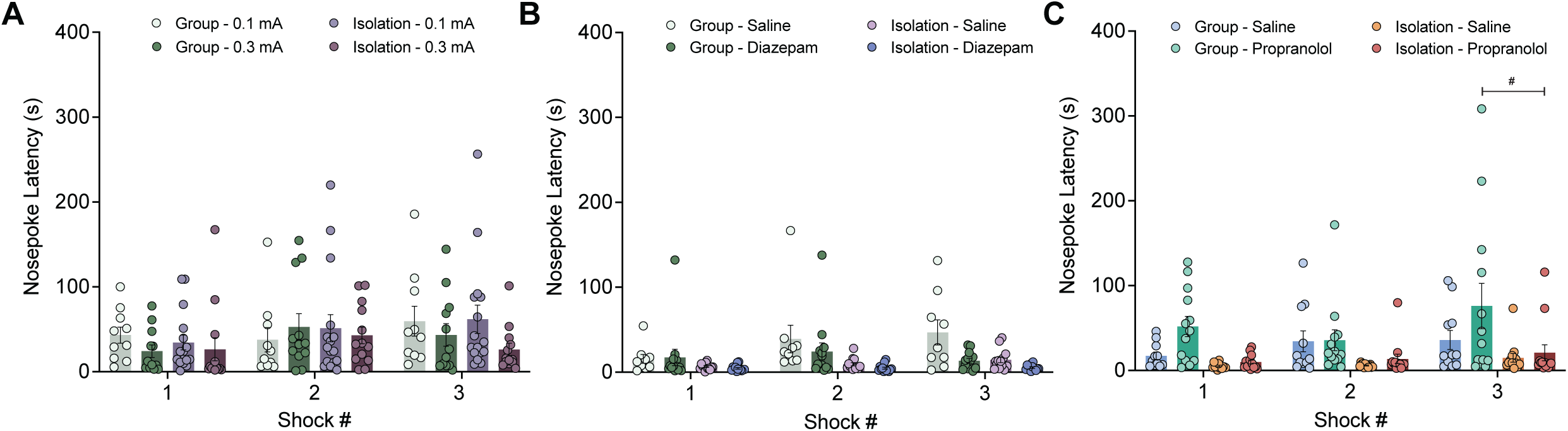
Footshock experience does not impact post-shock trial nosepoke latency. **(A)** Housing conditions and shock intensity have no effect on post-shock trial nosepoke latency **(B)** Housing conditions and diazepam (2 mg/kg) treatment have no effect on post-shock trial nosepoke latency. **(C)** Housing conditions and propranolol treatment (10 mg/kg) increase nosepoke latency for trials immediately following a shock trial; Group-Propranolol-Block 3 vs. Isolation-Propranolol-Block 3, # *p* < 0.05. Data are presented as mean ± SEM with dots reflecting individual subjects. For comparisons and exact p-values see **Table S1.**

## Notes

### Competing Interest Statement

The authors have declared no competing interest.

### Summary of Updates

The text has been edited to be more clear regarding the methods and tasks completed. Further analyses have been performed leading to new figure panels 1E, 2F-H, 3F-H. and 3N-P. As a result a new author has also been added.

## References

1. American Psychiatric Association, editor. Diagnostic and statistical manual of mental disorders: DSM-5. 5th ed. Washington, D.C: American Psychiatric Association; 2013.

2. Simmons JM, Winsky L, Zehr JL, Gordon JA. Priorities in stress research: a view from the U.S. National Institute of Mental Health. Stress. 2021;24:123–129.

3. Belzung C, Griebel G. Measuring normal and pathological anxiety-like behaviour in mice: a review. Behav Brain Res. 2001;125:141–149.

4. Bourin M. Animal models for screening anxiolytic-like drugs: a perspective. Dialogues Clin Neurosci. 2015;17:295–303.

5. Cryan JF, Sweeney FF. The age of anxiety: role of animal models of anxiolytic action in drug discovery: Age of anxiety. British Journal of Pharmacology. 2011;164:1129–1161.

6. Choleris E, Thomas a W, Kavaliers M, Prato FS. A detailed ethological analysis of the mouse open field test: effects of diazepam, chlordiazepoxide and an extremely low frequency pulsed magnetic field. Neuroscience and Biobehavioral Reviews. 2001;25:235– 260.

7. Irvine EE, Cheeta S, File SE. Tolerance to nicotine’s effects in the elevated plus-maze and increased anxiety during withdrawal. Pharmacol Biochem Behav. 2001;68:319–325.

8. Walf AA, Frye CA. The use of the elevated plus maze as an assay of anxiety-related behavior in rodents. Nat Protocols. 2007;2:322–328.

9. Shepherd JK, Grewal SS, Fletcher A, Bill DJ, Dourish CT. Behavioural and pharmacological characterisation of the elevated ‘zero-maze’ as an animal model of anxiety. Psychopharmacology (Berl). 1994;116:56–64.

10. La-Vu M, Tobias BC, Schuette PJ, Adhikari A. To Approach or Avoid: An Introductory Overview of the Study of Anxiety Using Rodent Assays. Frontiers in Behavioral Neuroscience. 2020;14.

11. Park J, Moghaddam B. Risk of punishment influences discrete and coordinated encoding of reward-guided actions by prefrontal cortex and VTA neurons. Elife. 2017;6.

12. Lezak KR, Missig G, Carlezon WA. Behavioral methods to study anxiety in rodents. Dialogues Clin Neurosci. 2017;19:181–191.

13. Dickson PE, Mittleman G. Strain and sex dependent effects of isolation housing relative to environmental enrichment on operant sensation seeking in mice. Sci Rep. 2021;11:17826.

14. Haluk DM, Wickman K. Evaluation of study design variables and their impact on food-maintained operant responding in mice. Behavioural Brain Research. 2010;207:394–401.

15. Arndt SS, Laarakker MC, Van Lith HA, Van Der Staay FJ, Gieling E, Salomons AR, et al. Individual housing of mice — Impact on behaviour and stress responses. Physiology & Behavior. 2009;97:385–393.

16. Võikar V, Polus A, Vasar E, Rauvala H. Long-term individual housing in C57BL/6J and DBA/2 mice: assessment of behavioral consequences: Effect of isolation on mouse behavior. Genes, Brain and Behavior. 2004;4:240–252.

17. McCall JG, Al-Hasani R, Siuda ER, Hong DY, Norris AJ, Ford CP, et al. CRH Engagement of the Locus Coeruleus Noradrenergic System Mediates Stress-Induced Anxiety. Neuron. 2015;87:605–620.

18. McCall JG, Siuda ER, Bhatti DL, Lawson LA, McElligott ZA, Stuber GD, et al. Locus coeruleus to basolateral amygdala noradrenergic projections promote anxiety-like behavior. ELife. 2017. https://elifesciences.org/articles/18247. Accessed 19 July 2018.

19. Faul F, Erdfelder E, Lang A-G, Buchner A. G*Power 3: a flexible statistical power analysis program for the social, behavioral, and biomedical sciences. Behav Res Methods. 2007;39:175–191.

20. Rivera-Irizarry JK, Skelly MJ, Pleil KE. Social Isolation Stress in Adolescence, but not Adulthood, Produces Hypersocial Behavior in Adult Male and Female C57BL/6J Mice. Front Behav Neurosci. 2020;14:129.

21. Granville-Grossman KL, Turner P. The effect of propranolol on anxiety. Lancet. 1966;1:788– 790.

22. Gorman a L, Dunn a J. Beta-adrenergic receptors are involved in stress-related behavioral changes. Pharmacology, Biochemistry, and Behavior. 1993;45:1–7.

23. Jacobs DS, Allen MC, Park J, Moghaddam B. Learning of probabilistic punishment as a model of anxiety produces changes in action but not punisher encoding in the dmPFC and VTA. ELife. 2022;11:e78912.

24. Chowdhury TG, Wallin-Miller KG, Rear AA, Park J, Diaz V, Simon NW, et al. Sex differences in reward-and punishment-guided actions. Cogn Affect Behav Neurosci. 2019;19:1404–1417.

25. Griebel G, Belzung C, Perrault G, Sanger DJ. Differences in anxiety-related behaviours and in sensitivity to diazepam in inbred and outbred strains of mice. Psychopharmacology 2000 148:2. 2000;148:164–170.

26. Pádua-Reis M, Nôga DA, Tort ABL, Blunder M. Diazepam causes sedative rather than anxiolytic effects in C57BL/6J mice. Sci Rep. 2021;11:9335.

27. Podhorna J, McCabe S, Brown RE. Male and female C57BL/6 mice respond differently to diazepam challenge in avoidance learning tasks. Pharmacology Biochemistry and Behavior. 2002;72:13–21.

28. An X-L, Zou J-X, Wu R-Y, Yang Y, Tai F-D, Zeng S-Y, et al. Strain and Sex Differences in Anxiety-Like and Social Behaviors in C57BL/6J and BALB/cJ Mice. Exp Anim. 2011;60:111–123.

29. Yokota S, Suzuki Y, Hamami K, Harada A, Komai S. Sex differences in avoidance behavior after perceiving potential risk in mice. Behav Brain Funct. 2017;13:9.

30. Zhou J, Hormigo S, Sajid MS, Castro-Alamancos MA. Caution Influences Avoidance and Approach Behaviors Differently. J Neurosci. 2022;42:5899–5915.

31. Sartori SB, Landgraf R, Singewald N. The clinical implications of mouse models of enhanced anxiety. Future Neurology. 2011;6:531–571.

32. Hur J, Smith JF, DeYoung KA, Anderson AS, Kuang J, Kim HC, et al. Anxiety and the Neurobiology of Temporally Uncertain Threat Anticipation. J Neurosci. 2020;40:7949– 7964.

33. Grupe DW, Nitschke JB. Uncertainty and anticipation in anxiety: an integrated neurobiological and psychological perspective. Nat Rev Neurosci. 2013;14:488–501.

34. Tye KM, Prakash R, Kim S-Y, Fenno LE, Grosenick L, Zarabi H, et al. Amygdala circuitry mediating reversible and bidirectional control of anxiety. Nature. 2011;471:358–362.

35. Jennings JH, Sparta DR, Stamatakis AM, Ung RL, Pleil KE, Kash TL, et al. Distinct extended amygdala circuits for divergent motivational states. Nature. 2013;496:224–228.

36. Kim S-Y, Adhikari A, Lee SY, Marshel JH, Kim CK, Mallory CS, et al. Diverging neural pathways assemble a behavioural state from separable features in anxiety. Nature. 2013;496:219–223.

37. Felix-Ortiz AC, Beyeler A, Seo C, Leppla CA, Wildes CP, Tye KM. BLA to vHPC inputs modulate anxiety-related behaviors. Neuron. 2013;79:658–664.

38. Felix-Ortiz AC, Burgos-Robles A, Bhagat ND, Leppla CA, Tye KM. Bidirectional modulation of anxiety-related and social behaviors by amygdala projections to the medial prefrontal cortex. Neuroscience. 2016;321:197–209.

39. Adhikari A, Lerner TN, Finkelstein J, Pak S, Jennings JH, Davidson TJ, et al. Basomedial amygdala mediates top-down control of anxiety and fear. Nature. 2015;527:179–185.

40. Adhikari A, Topiwala MA, Gordon JA. Single units in the medial prefrontal cortex with anxiety-related firing patterns are preferentially influenced by ventral hippocampal activity. Neuron. 2011;71:898–910.

41. Adhikari A, Topiwala MA, Gordon JA. Synchronized activity between the ventral hippocampus and the medial prefrontal cortex during anxiety. Neuron. 2010;65:257.

42. Anthony T, Dee N, Bernard A, Lerchner W, Heintz N, Anderson D. Control of Stress-Induced Persistent Anxiety by an Extra-Amygdala Septohypothalamic Circuit. Cell. 2014;156:522–536.

43. Kheirbek MA, Drew LJ, Burghardt NS, Costantini DO, Tannenholz L, Ahmari SE, et al. Differential control of learning and anxiety along the dorsoventral axis of the dentate gyrus. Neuron. 2013;77:955–968.

44. Heydendael W, Sengupta A, Beck S, Bhatnagar S. Optogenetic examination identifies a context-specific role for orexins/hypocretins in anxiety-related behavior. Physiology & Behavior. 2014;130:182–190.

45. Grillon C, Baas JP, Lissek S, Smith K, Milstein J. Anxious Responses to Predictable and Unpredictable Aversive Events. Behavioral Neuroscience. 2004;118:916–924.

46. Grillon C, Baas JMP, Pine DS, Lissek S, Lawley M, Ellis V, et al. The Benzodiazepine Alprazolam Dissociates Contextual Fear from Cued Fear in Humans as Assessed by Fear-potentiated Startle. Biological Psychiatry. 2006;60:760–766.

47. Schmitz A, Grillon C. Assessing fear and anxiety in humans using the threat of predictable and unpredictable aversive events (the NPU-threat test). Nat Protoc. 2012;7:527–532.

48. Vogel JR, Beer B, Clody DE. A simple and reliable conflict procedure for testing anti-anxiety agents. Psychopharmacologia. 1971;21:1–7.

49. Fischer BD, Licata SC, Edwankar RV, Wang Z-J, Huang S, He X, et al. Anxiolytic-like effects of 8-acetylene imidazobenzodiazepines in a rhesus monkey conflict procedure. Neuropharmacology. 2010;59:612–618.

50. Miles L, Davis M, Walker D. Phasic and Sustained Fear are Pharmacologically Dissociable in Rats. Neuropsychopharmacol. 2011;36:1563–1574.

